# Climate adaptation and vulnerability of foundation species in a global change hotspot

**DOI:** 10.1101/2022.03.30.486132

**Authors:** Cristóbal Gallegos, Kathryn A. Hodgins, Keyne Monro

**Author notes:** Joint senior authors (Kathryn A. Hodgins, Keyne Monro). Corresponding author: Cristóbal Gallegos.

## Abstract

Climate change is altering species ranges, and abundances within ranges, as populations become differentially adapted and vulnerable to the climates they face. Hence, characterising current ranges, whether species harbour and exchange adaptive genetic variants, and how variants are distributed across landscapes undergoing rapid change, is crucial to predicting responses to future climates and informing conservation strategies. Such insights are nonetheless lacking for most species of conservation concern. We characterise genomic patterns of neutral variation, climate adaptation, and climate vulnerability (the amount of genomic change needed to track climate change by adaptation) in sister foundation species, the endemic marine tubeworms *Galeolaria caespitosa* and *Galeolaria gemineoa*, across a sentinel region for climate change impacts. First, species are shown to be partly sympatric despite previous support for non-overlapping ranges, and genetically isolated despite known capacity for hybrid crosses to yield viable early offspring. Second, species show signals of polygenic adaptation, but to differing components of temperature and involving mostly different loci. Last, species are predicted to be differentially vulnerable to climate change, with *G. gemineoa* — the less genetically diverse species — needing double the adaptation to track projected changes in temperature compared to its sister species. Together, our findings provide new insights into climate adaptation and its potential disruption by climate change for foundation species that enhance local biodiversity, with implications for evolutionarily-enlightened management of coastal ecosystems.

## Introduction

Global climate change is redistributing Earth’s biodiversity. Geographic ranges are shifting as species move to track tolerable climatic conditions, and abundances are changing within ranges as populations adapt, or grow maladapted and thereby vulnerable, to the climates they face (Pecl et al., 2017; Scheffers et al., 2016). Understanding current ranges, whether species harbour (and exchange) different genetic variants involved in climate adaptation, and how such variants are distributed across landscapes undergoing rapid climate change, is therefore key to predicting responses to future change and informing conservation strategies (Teixeira & Huber, 2021; Willi et al., 2022). This remains challenging for many species, especially those that are cryptic or unsuited to traditional ways of inferring adaptation and persistence (reciprocal transplants, multi-generation breeding experiments, etc.). However, emerging tools linked to the rise of population genomics for non-model organisms in recent years are set to provide new insights into climate adaptation and vulnerability for understudied species of conservation concern (Hoffmann et al., 2021; Hohenlohe et al., 2021).

Genomic prediction of climate adaptation relies on genome scans and genotype-environment associations to identify putatively adaptive loci harbouring variants (alleles) whose frequencies covary with climate across species ranges (Forester et al., 2016; Rellstab et al., 2015). Then, using machine learning-or distance-based methods and climate forecasts, climate-adaptive variants can be projected across space and through time to assess genomic vulnerability (also called genetic offset) as the predicted difference in their distributions across present and future landscapes (Fitzpatrick & Keller, 2015) — in other words, the amount of genomic change needed to track climate change via evolutionary adaptation (Capblancq et al., 2020; Hoffmann et al., 2021). Notwithstanding the challenges of validating predictions (Hoffmann et al., 2021; Rellstab et al., 2021), assessing genomic vulnerability offers new scope to ask how populations and species of high ecological importance, but limited tractability to experimentation, may fare in future climates, identifying those at most risk of decline as those needing to evolve the most to keep pace with change and avert maladaptation. Combining such assessments with insights from neutral genomic variation, moreover, allows population structures and species barriers to be explored from both neutral and adaptive perspectives, with differing implications for population dynamics, species range shifts, and management actions under climate change (Hohenlohe et al., 2021; Kardos et al., 2021; Willi et al., 2022).

Accordingly, mounting studies have assessed genomic vulnerability in the context of climate change for individual species — mostly trees (Borrell et al., 2020; Ingvarsson & Bernhardsson, 2020; Jia et al., 2020; Pina-Martins et al., 2019) or marine counterparts (Vranken et al., 2021; Wood et al., 2021), but also birds (Bay et al., 2018). Yet rarely, if ever, has the approach been extended to related species in overlapping ranges (but see Nielsen et al., 2021), despite the impacts of dispersal and gene flow not just across populations, but across partial species barriers. Introducing new adaptive variants from one population or species to another, for example, may create highly-fit hybrids that increase population sizes in the short term (Fitzpatrick et al., 2020) or rates of adaptation in the longer term (Grant & Grant, 2019; Mitchell et al., 2019). Conversely, it may cause outbreeding depression if distantly-related genomes are less compatible (Frankham, 2015), or expose variants to new environments in which they are maladapted (Hoffmann & Sgrò, 2011; Polechová, 2018). Over time, species lines may blur, or species that are less vulnerable to climate change may displace species that are more so, at a net cost to biodiversity (Román-Palacios & Wiens, 2020; Todesco et al., 2016). From this perspective, multi-species assessments of genomic vulnerability may help to identify whether genetic lineages are on distinct (and potentially adaptive) evolutionary pathways linked to climate, and could therefore warrant separate management to conserve their genetic uniqueness (Willi et al., 2022).

Gaps also exist in our understanding of adaptation and vulnerability to different components of climate change, which is altering not only the mean values (trends) of key variables, but also their variability, extremes, and the extents to which they vary predictably or stochastically (Fischer & Knutti, 2015; Ruokolainen et al., 2009; Waldock et al., 2018). By imposing different selective pressures, these components of climate change may have different consequences for biodiversity and lead to different risks of population decline (Bitter et al., 2021; Kingsolver & Buckley, 2017; Lande, 2014; Rescan et al., 2021; Ripa & Lundberg, 1996). To date, however, most assessments focus on adaptation to climate variables or proxies (precipitation, temperature, vegetation, elevation) relevant to terrestrial systems, whereas marine systems are underrepresented by comparison (Grummer et al., 2019; Lotterhos et al., 2021). Marine species often have high fecundity, large effective population sizes, and long-range dispersal at early life stages (gametes, embryos, and larvae) with high mortality, so that gene flow, selection, and drift play out in oceanographic settings that can strongly couple physical and evolutionary processes, while also decoupling the environments of early stages and adults. Trends in key variables (such as temperature), moreover, are less striking and immutable than they are on land (Gaylord & Gaines, 2000), potentially giving other components of change greater influence. Marine systems can therefore offer new genomic insights into climate adaptation and vulnerability (Liggins et al. 2020), but studies remain rare (Vranken et al., 2021; Wood et al., 2021). They have not explored adaptation to environmental predictability, and are lacking for many species of ecological importance in regions undergoing rapid climate change where increased adaptation can be expected (Hill et al., 2011; Lotterhos et al., 2021).

Southeast Australia is a climate change and biodiversity hotspot, identified as one of the world’s fastest warming marine regions and one of its most biologically diverse (Frusher et al., 2014; Hobday & Pecl, 2014; Ramírez et al., 2017). East-west divergence of populations and species in the region is often attributed to geographic isolation by the historical land-bridge joining Tasmania and mainland Australia during the last glacial maxima (Dawson, 2005; O’Hara & Poore, 2000). The region also sees two boundary currents — the East Australian Current flowing south from the tropics, and the Zeehan Current flowing east from the Great Australian Bight — converge with subantarctic water in Bass Strait, generating complex gradients of temperature and flow that may mediate postglacial dispersal, drift, and selection (Miller et al., 2020; Waters, 2008). Those gradients are set to steepen as the East Australian Current continues to warm and intensify southward (Hobday & Lough, 2011; Ridgway & Hill, 2009), making the region a natural laboratory for studying climate adaptation and vulnerability in order to better predict the fate of biodiversity in future climates.

Here, we investigate climate adaptation and vulnerability in an endemic ecosystem engineer, or foundation species — the marine tubeworm, *Galeolaria* — across the southeast hotspot. *Galeolaria* comprises cryptic sister species that are geographically concordant with neutral genetic markers (placing *G. gemineoa* to the northeast and *G. caespitosa* to the southwest; Halt et al., 2009) yet are still able to interbreed (Styan et al., 2008). Their ranges, population structures, frequency of hybridization, and potential adaptation to climate are unknown. We therefore characterized genomic variation among populations of each species throughout the hotspot to assess genomic divergence, diversity, and gene flow within and between species. We further identified candidate adaptive loci and associations with different components of temperature for each species, then modelled allele turnover at candidate loci in current and projected climates to predict where populations are most vulnerable to loss of adaptation with ongoing climate change. Our analyses reveal these species to be genetically distinct despite partial sympatry across the hotspot, support climate adaptation in both species, and identify populations that could face greater risk of decline unless they adapt rapidly to near-future climates. Such insights into the nature of biodiversity across the hotspot could enhance evolutionarily-enlightened management and conservation strategies in a sentinel region for understanding climate impacts.

## Methods

### Study system

*Galeolaria* is an ecosystem engineer endemic to rocky shores of southeast Australia, where its dense colonies of stony tubes enhance local biodiversity by providing habitat and climate refugia for species that cannot otherwise persist there (Figure 1A; Wright & Gribben, 2017). Year-round, adults release gametes into the sea for external fertilization and embryogenesis (Chirgwin et al., 2020, 2021), then larvae spend ~2–3 weeks offshore, dispersed by currents, before transitioning to sessile life stages (juveniles and adults) onshore in the intertidal. As for other aquatic ectotherms, planktonic stages are thermal bottlenecks in the lifecycle, defining vulnerability to climate as well as population structure across species’ ranges (Dahlke et al., 2020; Lotterhos et al., 2021; Rebolledo et al., 2020). *Galeolaria caespitosa* and *G. gemineoa* are said to diverge in the southeast hotspot near Ninety Mile Beach, due to historical vicariance, dispersal limitation, or lack of rocky habitat (Figure 1B; Styan et al. 2008; Halt et al. 2009).

**Figure 1.**
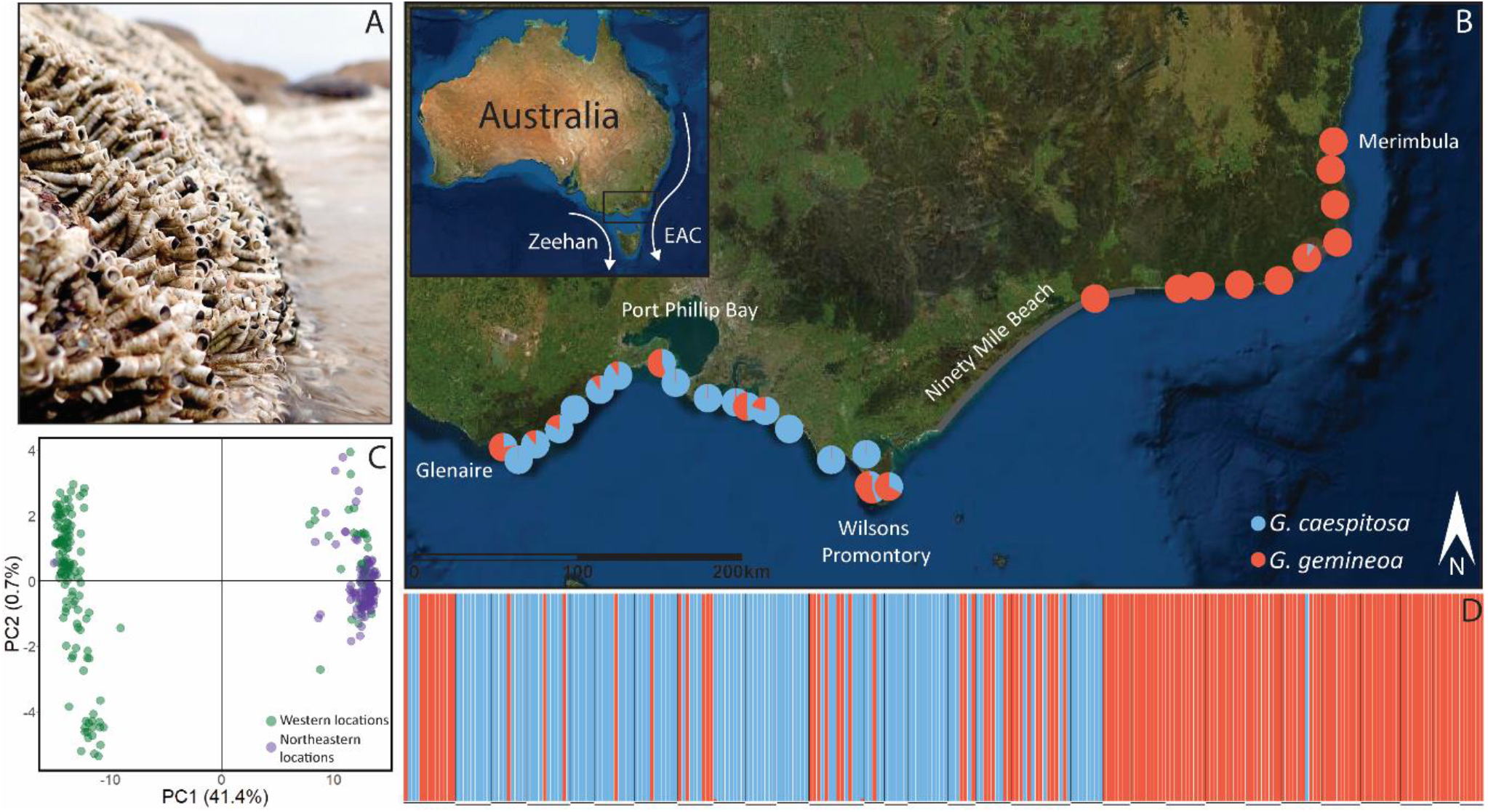
Geographic setting and genetic structure of *Galeolaria*. (A) A typical colony showing adults retracted into tubes at low tide. (B) Locations from which individuals were sampled across the southeast hotspot, where boundary currents converge at a now-submerged land bridge between Tasmania and mainland Australia (inset). Pie charts show the proportions of individuals identified as *G. caespitosa* (blue) and *G. gemineoa* (red) by ADMIXTURE analyses. Until now, species ranges were thought to diverge near Ninety Mile Beach (grey line), which lacks rocky habitat to colonise. (C) A principal components analysis of genetic variation reveals two distinct clusters corresponding to the two species, with individuals from western locations (Glenaire to Wilsons Promontory) in green and individuals from northeastern locations (Wilsons Promontory to Merimbula) in purple. (D) Ancestries of individuals (vertical bars, coloured as in panel A) suggest little gene flow between species. Horizontal lines below bars group individuals by location.

### Sampling throughout the southeast Australian hotspot

We sampled adult populations of *G. caespitosa* and *G. gemineoa* from 30 locations spanning ~800 km of coast throughout the hotspot (Figure 1B; Table S1) in January 2019. Locations were separated by ~20 km (subject to accessibility and species detection) and were chosen to capture thermal variation in each species’ range. Each of 10 to 15 individuals per location was immediately extracted from its tube, spawned for 5 minutes in filtered seawater to minimize contamination by gametes, then rinsed and placed in an Eppendorf tube with 70% ethanol. Individuals were transported to the lab and stored at room temperature (~22 °C) until DNA extraction.

### DNA extraction, library preparation, and sequencing

We extracted DNA from the posterior ~5 mm of each individual. We digested tissue overnight with proteinase K, then extracted DNA using the Qiagen DNeasy Blood and Tissue Kit following manufacturer instructions (Qiagen, 2006). Quality was checked by running individual samples on 2% agarose gel stained with ethidium bromide and also with a UV-Vis Spectrometer (NanoDrop 1000, Thermo Scientific). Quantity was checked with a QuBit fluorometer (dsDNA HS, Invitrogen).

Library preparation followed a double-digest (with *HF-PstI* and *MspI* restriction enzymes), genotype-by-sequencing (ddGBS) protocol with equal amounts of DNA per individual (Poland et al., 2012). The protocol was modified by performing PCR reactions for individual samples, then pooling equal amounts of the PCR products. A size selection step was also added to focus on fragments between 400-600 bp. Sequencing was performed in two batches, one by GenomeQuébec (Montréal, Canada) and one by GENEWIZ (Suzhou, China). Both batches used one lane of Illumina HiSeq 4000 (paired-end, 150 bp).

### Identifyng single nucleotide polymorphisms (SNPs)

Reads were quality-checked using FastQC (https://www.bioinformatics.babraham.ac.uk/projects/fastqc/), then demultiplexed and cleaned using *process_radtags* in the Stacks software pipeline (Catchen et al., 2013; Catchen et al., 2011). To optimise parameter values for identifying SNPs, we ran the pipeline nine times using a range of values for a subset of 16 individuals, then explored key statistics including the distribution of SNPs per locus and the number of loci shared by at least 80% of individuals (Paris, Stevens, and Catchen 2017; Rochette and Catchen 2017). Based on results, we called SNPs for all individuals using values of *m* = 3, *M* = 5, and *n* = 5.

We filtered SNPs in several steps. First, we filtered loci with low allele frequencies in *Stacks* (with *min_maf* = 0.01). Next, we filtered the remainder in *vcfR* v1.8.0 (Knaus & Grünwald, 2017), *adegenet* v2.1.1 (Jombart, 2008), and gaston v1.5.6 (Perdry & Dandine-Roulland, 2020) to keep only biallelic loci with depth > 5, genotype quality > 30, and linkage disequilibrium (r^2^) < 0.8, and to exclude individuals missing more than 60% of loci. Last, we excluded loci missing more than 55% of data across individuals and used this large SNP set for genotype-environment association analyses. For analyses of population genetic structure, which do not require large numbers of SNPs, we used a reduced SNP set that excluded loci missing more than 30% of data across individuals.

All remaining data analyses were performed in R v.4.0.5 (R Core Team, 2021) unless otherwise stated.

### Environmental data

We obtained a high-resolution (1 km^2^ grid cell) satellite-based time series of sea surface temperature from January 2010 to December 2018 (www.ghrsst.org), and extracted daily observations for all 30 locations. We summarised data using ten variables based mostly on the WorldClim scheme (Fick & Hijmans, 2017; Hijmans et al., 2005), then selected four that captured different components of change in temperature and had pairwise correlations below |0.7| (Table S2). They were the mean temperature (a measure of trend), maximum temperature of the warmest month (a measure of extremity), mean monthly temperature range (a measure of variability), and temperature noise structure (a measure of stochasticity). See supplementary material for details of calculations.

### Analyses of population genetic structure

#### Genetic clustering

To explore genetic differentiation between populations of each species, we clustered loci using a principal components analysis of genetic variation in the *adegenet* package (v2.1.1; Jombart, 2008). We also estimated the ancestries of individuals, and levels of admixture among ancestral lineages, using the ADMIXTURE program (Alexander et al., 2009). The parameter *K* (presumed number of ancestral lineages) was set to 2, which minimized cross-validation error in preliminary analyses. Setting higher values of *K* did not alter our results.

### Genetic diversity

To explore genetic diversity within populations of each species, and because some loci were polymorphic between species but not within them, we filtered out loci that were monomorphic or unique to one species. For each population, we then calculated standard measures of diversity — observed heterozygosity (H_O_), expected heterozygosity (H_S_), inbreeding coefficient (F_IS_), and allelic richness (AR) — averaged across loci in *hierfstat* v0.5-7 (Goudet & Jombart, 2015). We compared diversity between species using *F*-tests from linear models with species as a categorical fixed effect. Checks of model assumptions using diagnostic plots of residuals detected no serious violations.

### Genetic isolation by distance

To identify evidence for greater gene flow among geographically proximate populations of each species, we first calculated pairwise genetic distances (F_ST_, also averaged across loci) between populations in *hierfstat* v0.5-7 (Goudet & Jombart, 2015). To reduce sampling error, populations represented by less than three individuals were excluded from calculations (Nazareno et al., 2017). To assess the relationship between genetic isolation and geographic distance in each species, we tested the correlation between matrices of pairwise genetic distances (F_ST_/1 – F_ST_) and geographic distances (calculated with the *geosphere* package; (Hijmans et al., 2017) using a Mantel test based on 999 permutations in *dartR* v1.8.3 (Gruber & Georges, 2019).

### Genetic isolation by temperature

To assess the relationship between genetic isolation and thermal environment in each species, we calculated environmental distances based on temperature variables in the *ade4* package (Dray & Dufour, 2007), then tested their correlation with genetic distances using a Mantel test in the same package.

### Analyses of climate adaptation and vulnerability

#### Candidate loci for thermal adaptation

To search for candidate adaptive loci that diverge among populations in association with temperature variables, we performed a redundancy analysis for each species using the *vegan* package (Oksanen et al., 2016). This two-step extension of linear regression to multivariate responses identifies loci that covary in response to multivariate environments, and provides a superior combination of low false-positive and high true-positive rates to other methods (Forester et al., 2018). Here, it involved regressing loci on temperature variables to compute a matrix of predicted genotype-temperature associations, then applying principal components analysis to the matrix to compute four uncorrelated principal components, or ordination axes (RDA1– RDA4), comprising linear combinations of variables that explain those associations. Candidate loci were identified as outliers on ordination axes based on scores at least three standard deviations from the mean score per axis (two-tailed *P*-value = 0.003). Because the method does not tolerate missing data, missing genotypes were imputed by population using the most common genotype per locus. If loci were missing or tied, we used the most common genotype in all samples (Forester et al., 2018).

To cross-validate results with those from alternative approaches, we performed equivalent univariate analyses using the standard covariate model and default settings in the BayPass program (Gautier, 2015). We identified candidate loci based on *P*-values of XtX statistics (F_ST_ analogues accounting for population structure) and inferred associations with temperature variables based on Bayes factors greater than ten (“strong evidence”; Gautier 2015). We then tested the overlap of candidates identified by each method using one-tailed hypergeometric tests. As further cross-validation, we repeated the redundancy analysis with distance-based Moran’s eigenvector maps included to account for population structure (Forester et al., 2018).

#### Genomic vulnerability to climate change

To predict each species’ vulnerability to future climate change, we modelled temperature-driven turnover in alleles at candidate loci using gradient forest regression models (Fitzpatrick & Keller, 2015), then mapped current and future turnovers throughout the study range. We fitted each model using minor allele frequencies at candidate loci that overlapped redundancy and BayPass analyses as the response variables, temperature variables as predictors, and constructed 2000 regression trees per locus using default settings in the *gradientForest* package (Ellis et al., 2012).

To map current turnover, we extracted temperature variables for each grid cell in the study range and transformed variables into genetic importance (relative contributions to turnover) using the turnover function estimated by the model (Fitzpatrick & Keller, 2015). We then summarised transformed variables as three principal components, assigned each component to a RGB colour palette following Ellis et al. (2012), and mapped colours to grid cells using the *raster* package (Hijmans, 2017). Mapped this way, colours predict genetic compositions (allele frequencies) in cells, and locations with similar colours are predicted to harbour populations with similar compositions. Biplots of the two largest principal components were used to relate turnover in composition to changes in temperature (Ellis et al., 2012).

Rather than map future turnover directly, we translated it to the genetic offset needed to maintain thermal adaptation under climate change (Ellis et al., 2012; Fitzpatrick & Keller, 2015). To do so, we repeated the process above with mean and maximum temperatures projected for 2050 and 2100 under low (RCP45) and high (RCP85) CO_2_ emission scenarios, extracted for each grid cell from the Bio-ORACLE database (Assis et al., 2018; Tyberghein et al., 2012). Since other variables were unavailable, we also re-calculated current turnover without them. For each cell, we transformed variables into genetic importance as above, calculated genetic offset as the Euclidian distance between current and future genetic compositions, then mapped offset as above.

## Results

### Variant identification

Sequencing returned an average of 3,392,340 quality-filtered reads per individual, with an average depth of coverage of 24.1-fold. Stacks (Catchen et al., 2011) identified 8,887,109 putative SNPs from 330 individuals. Filtering retained 8,788 unlinked loci from 272 individuals (with 16.1% of data missing across loci and individuals) for analyses of population genetic structure, and 24,263 unlinked loci from 272 individuals (with 31.3% of data missing across loci and individuals) for genotype-environment association analyses.

### Analyses of population genetic structure

#### Genetic clustering

Principal components analysis of genetic variation revealed two distinct genetic clusters defined by PC1 and PC2, jointly accounting for 42.1% of the multilocus genetic variation sampled (Figure 1C). Most individuals in one cluster were sampled northeast of Wilsons Promontory (to Merimbula, our northernmost location; Figure 1B), whereas most individuals in the other cluster were sampled west of this point (to Glenaire, our westernmost location; Figure 1B). However, multiple individuals from western locations clustered with the ‘northeastern’ cluster, and one individual from a northeastern location clustered with the ‘western’ cluster (Figure 1C).

ADMIXTURE analyses confirmed the presence of individuals from different ancestral lineages in western and northeastern populations (Figure 1B and 1D), but detected little gene flow between lineages (individual ancestry proportions consistently exceeded 0.99, shown by single-coloured bars in Figure 1D). Based on known distributions of *Galeolaria* species (Halt et al., 2009), the ‘northeastern’ cluster is *G. gemineoa* and the ‘western’ cluster is *G. caespitosa*, but species are now shown to be sympatric in some locations, especially west of Wilsons Promontory (Figure 1B). Subsequent analyses were therefore separated by species. We also explored genetic clustering within species but detected none at this level (Figure S1), as was further confirmed by within-species ADMIXTURE analyses.

### Genetic diversity

Of the reduced SNP set, 2,495 loci were polymorphic within species and shared between species. On average, measures of genetic diversity based on these loci were significantly lower for *G. gemineoa* than for *G. caespitosa*, except for inbreeding coefficients (Table 1). These coefficients were positive and similar in magnitude (~0.21) for both species, indicating that their populations harbour fewer heterozygotes than expected under Hardy-Weinberg equilibrium.

**Table 1.**
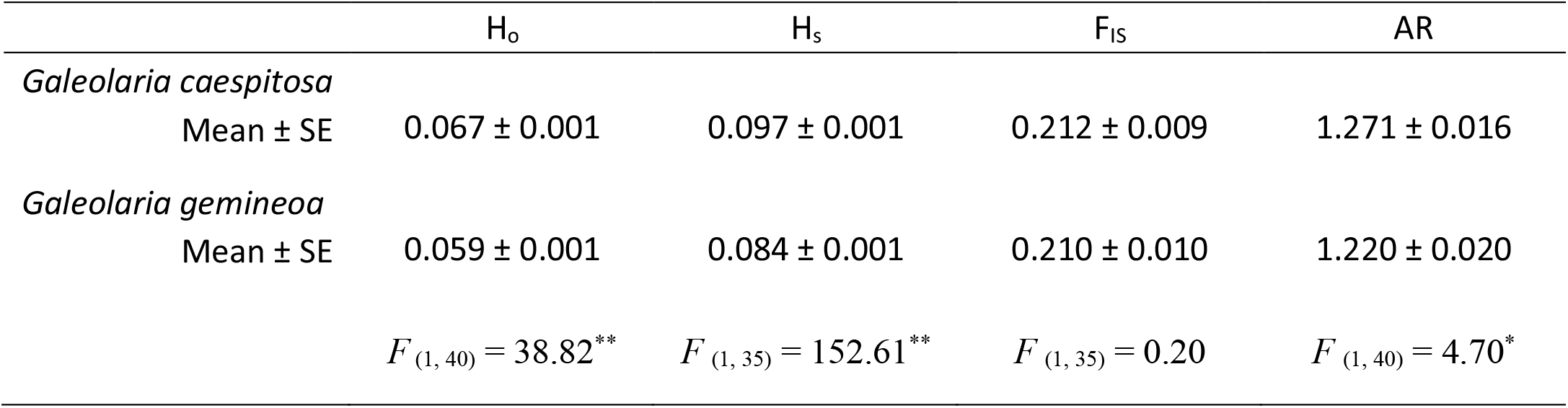
Estimates of genetic diversity for *G. caespitosa* and *G. gemineoa*. H_o_ is observed heterozygosity, H_S_ is expected heterozygosity, F_IS_ is the inbreeding coefficient, and AR is allelic richness (ranging from one to two because only biallelic loci were analysed). Estimates are averaged across loci and populations (see supplementary Table S3 for population values) and compared between species using *F*-tests (^*^*P* < 0.05; ^**^*P* < 0.001).

| | $H_o$ | $H_s$ | $F_{IS}$ | AR |
| --- | --- | --- | --- | --- |
| *Galeolaria caespitosa* |  |  |  |  |
| Mean ± SE | 0.067 ± 0.001 | 0.097 ± 0.001 | 0.212 ± 0.009 | 1.271 ± 0.016 |
| *Galeolaria gemineoa* |  |  |  |  |
| Mean ± SE | 0.059 ± 0.001 | 0.084 ± 0.001 | 0.210 ± 0.010 | 1.220 ± 0.020 |
| | $F_{(1, 40)} = 38.82^{**}$ | $F_{(1, 35)} = 152.61^{**}$ | $F_{(1, 35)} = 0.20$ | $F_{(1, 40)} = 4.70^*$ |

### Genetic isolation by distance

Mean pairwise genetic distance (F_ST_ ± SE; see Figure S2 for all values) was relatively low for both species (*G. caespitosa*: 0.066 ± 0.001; *G. gemineoa*: 0.062 ± 0.001) but significantly lower for *G. gemineoa* (*F* _(1, 305)_ = 6.567, *P* < 0.02). Mantel tests failed to detect an association between pairwise genetic distance and geographic distance for either species (*G. caespitosa*: *r* = 0.006, *P* = 0.508; *G. gemineoa*: *r* = 0.212, *P* = 0.058).

To further check whether species remain genetically isolated in sympatry, we compared mean species-level F_ST_ between sympatric and allopatric populations (Figure S3). No difference was detected (F_ST_ in sympatry: 0.599 ± 0.009; F_ST_ in allopatry: 0.598 ± 0.001; *F* _(1, 85)_ = 0.015, *P* = 0.903), suggesting that species barriers persist even when geographical barriers are absent.

#### Genetic isolation by temperature

Mantel tests also failed to detect an association between pairwise genetic distance and distance in thermal environment for either species (*G. caespitosa*: *r* = −0.119, *P* = 0.692; *G. gemineoa*: *r* = 0.003, *P* = 0.462).

### Analyses of climate adaptation and vulnerability

#### Candidate loci for thermal adaptation

Redundancy analyses identified significant associations between individual loci and temperature variables that explained ~2% of multilocus genetic variation and supported thermal adaptation in each species (*G. caespitosa*: adjusted *R*^2^ of global model = 0.021, *P* < 0.002; *G. gemineoa*: adjusted *R*^2^ of global model = 0.020, *P* < 0.002). Multiple, independent associations were inferred by the significance of all four ordination axes per analysis (*P <* 0.002), with the two largest axes jointly capturing 53% of associations detected in *G. caespitosa* and 56% of associations detected in *G. gemineoa* (Figure 2). Of 775 candidate loci detected in *G. caespitosa* (from 15,636 loci screened), 181 were most associated with mean temperature, 230 were most associated with maximum temperature, 213 were most associated with monthly temperature range, and 151 were most associated with temperature noise structure (Table S4). Association strengths (measured as correlations) averaged 0.354 and ranged from 0.077 to 0.769. Of 679 candidate loci detected in *G. gemineoa* (from 15,462 loci screened), 248 were most associated with mean temperature, 93 were most associated with maximum temperature, 173 were most associated with monthly temperature range, and 165 were most associated with temperature noise structure (Figure 2; Table S4). Association strengths averaged 0.311 and ranged from 0.081 to 0.856.

**Figure 2.**
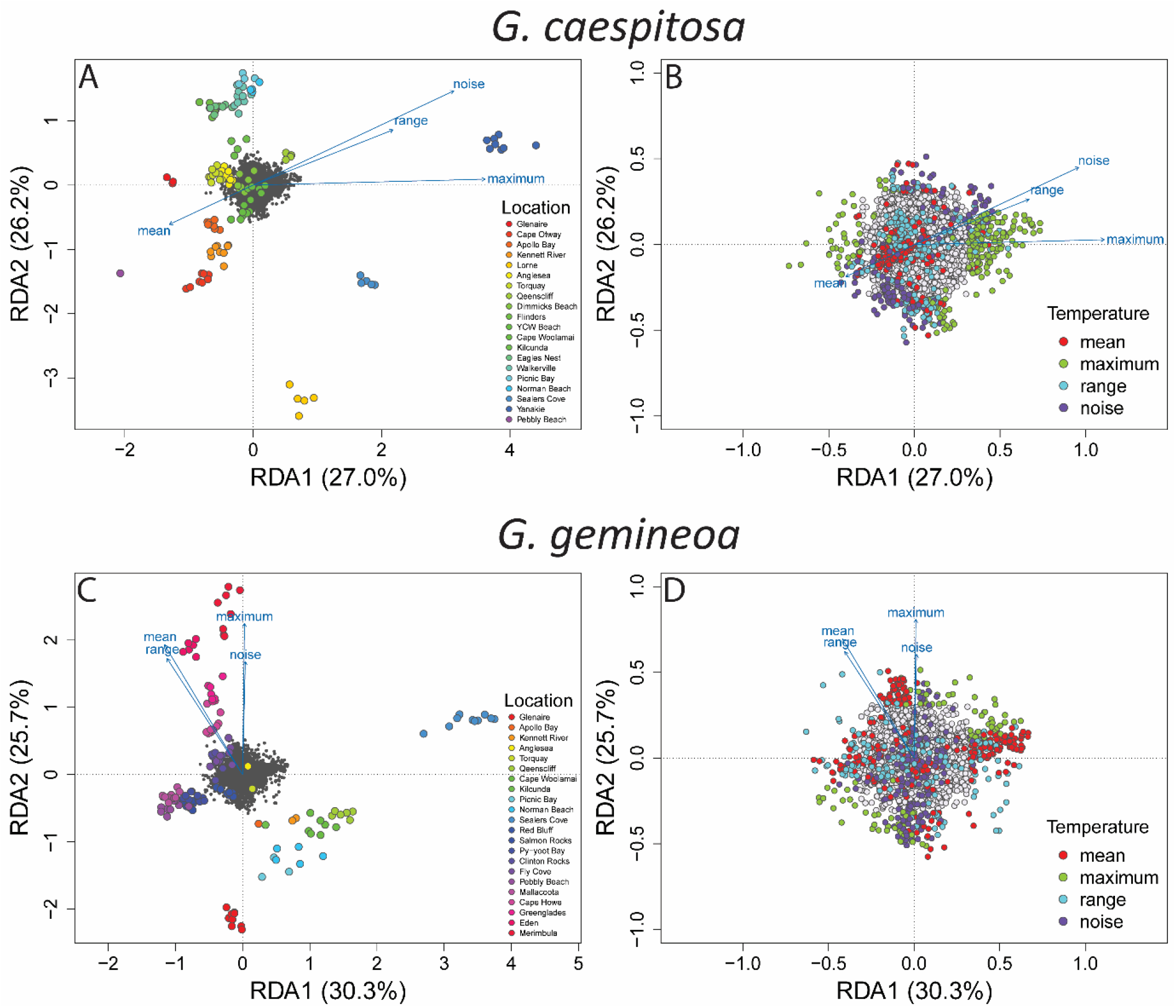
Associations between genotype and temperature identified for *G. caespitosa* (A–B) and *G. gemineoa* (C–D) by redundancy analysis. Biplots show the two largest ordination axes (RDA1 and RDA2) per analysis, comprising linear combinations of temperature variables (mean, maximum, range, and noise structure) that explain 53–56% of associations with multilocus genetic variation per species (see Figure S4 for other axes). In all panels, closer alignments of items with ordination axes indicate stronger associations with axes. In (A) and (C), grey points are single loci, other points are individuals coloured by location, and vectors are variables. In (B) and (D), which magnify left-hand plots to focus on loci, candidate adaptive loci (identified as significant outliers on ordination axes) are coloured by the variables they associate most strongly with.

BayPass analyses also identified multiple candidate loci associated with temperature variables, some of which overlapped those identified by redundancy analyses (Figure 3, top row), and did so for specific variables (Table S4). Robust candidates identified by both methods were used to further predict genomic vulnerability (see Figures 4 and 5). Few candidates overlapped between species (Figure 3, bottom row), which seemingly adapt to temperature using mostly different loci. Overlaps were generally higher than expected by chance (*P* < 0.001), except for the overlap between species resulting from redundancy analyses (Figure 3, bottom row).

**Figure 3.**
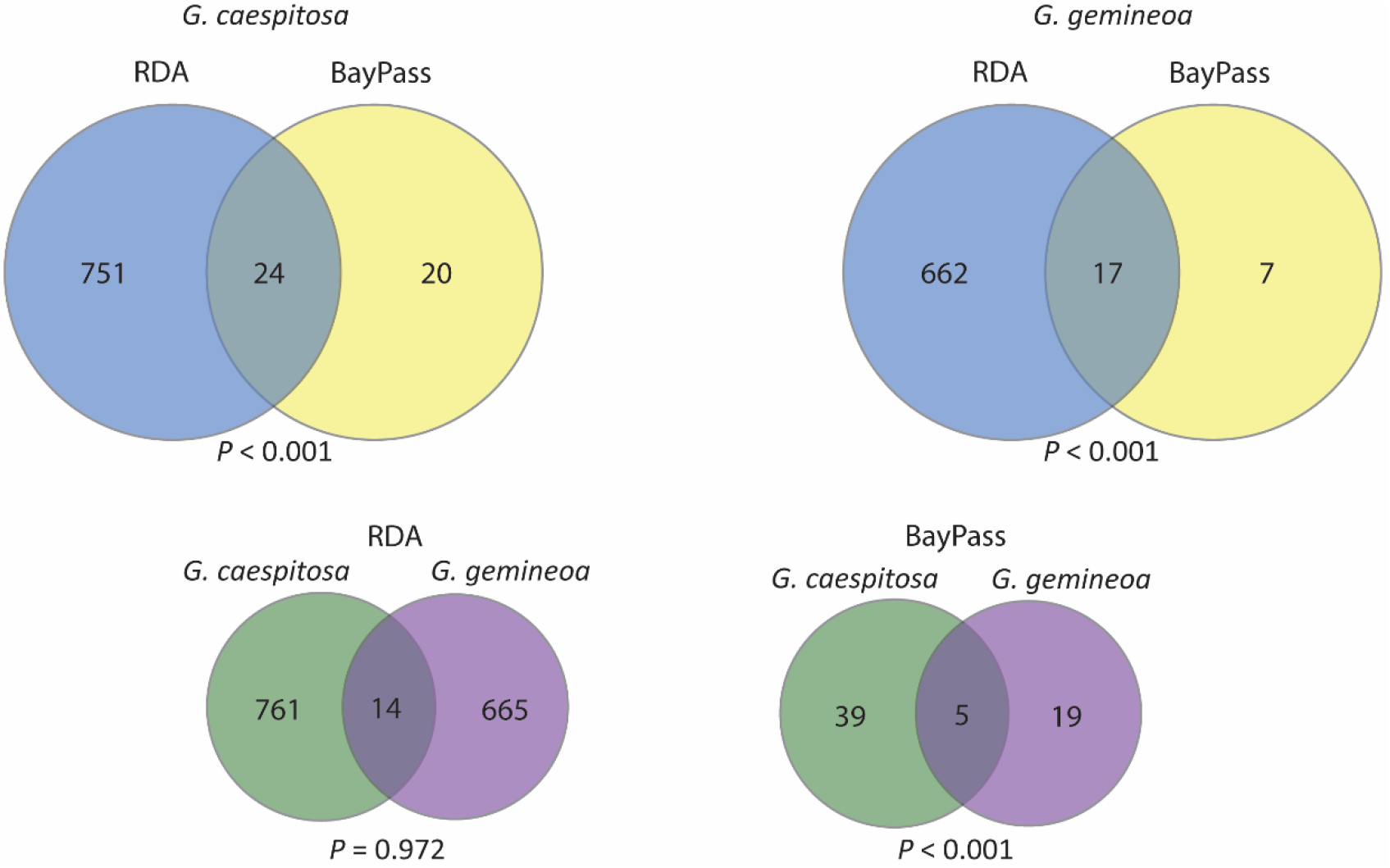
Candidate adaptive loci identified for *G. caespitosa* and *G. gemineoa* by redundancy analyses (RDA) *versus* BayPass analyses. The top row shows overlaps between methods for each species (overlapping candidates were used to further predict genomic vulnerability; see Figures 4 and 5). The bottom row shows overlaps between species for each method, suggesting that the genetic basis of adaptation mostly differs between species. *P*-values are the probabilities of observing overlaps by chance, given the numbers of candidates identified from the numbers of loci screened.

**Figure 4.**
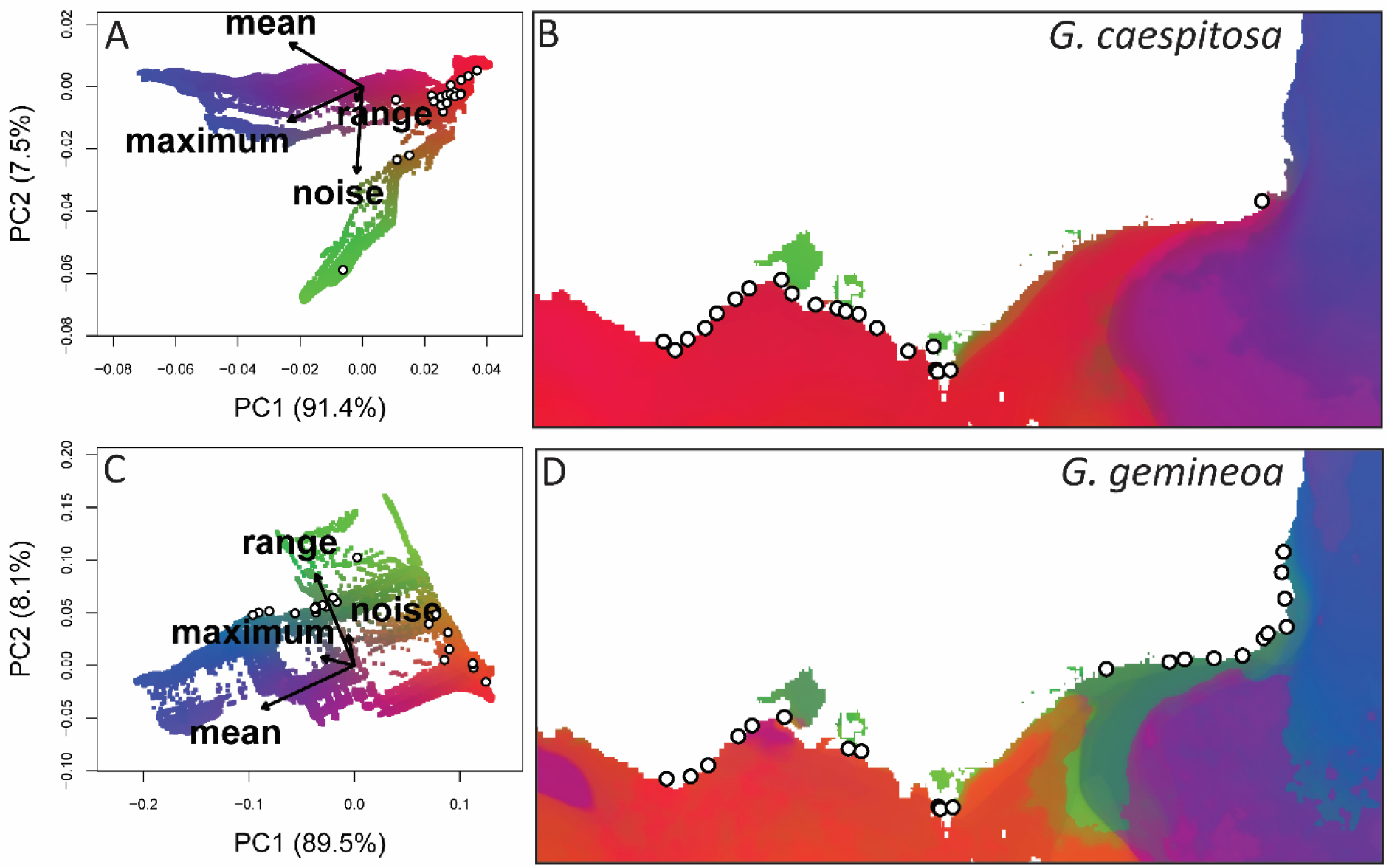
Temperature-driven turnover in alleles at candidate loci predicted for *G. caespitosa* (A–B) and *G. gemineoa* (C–D) by gradient forest models. Biplots in (A) and (C) show the two largest principal components (PC1 and PC2) per model, comprising linear combinations of temperature variables (mean, maximum, range, and noise colour) that explain 98–99% of allele turnover per species. Colours predict genetic compositions (allele frequencies) along biplot axes, and vectors relate compositions to variables (variables have higher values in the directions of vectors and lower values in opposing directions). Maps in (B) and (D) predict genetic compositions throughout the study range, and locations with similar colours are predicted to harbour populations with similar compositions. In all panels, points are locations from which individuals were sampled. Note that species have planktonic life stages (gametes, embryos, and larvae) that spend days to weeks offshore before transitioning to sessile life stages (juveniles and adults) onshore in the intertidal.

**Figure 5.**
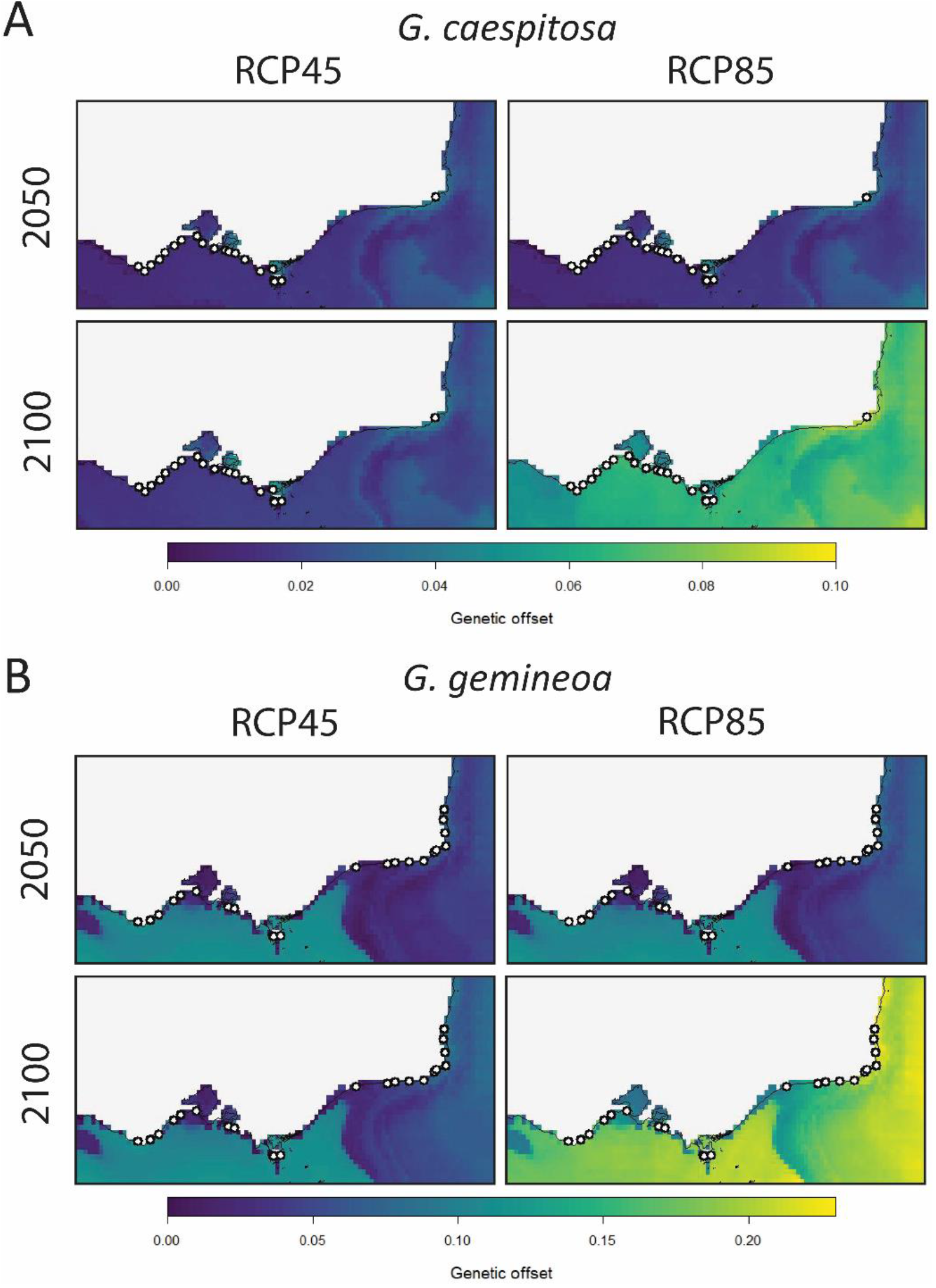
Genetic offsets needed to maintain thermal adaptation under future climate change for *G. caespitosa* (A) and *G. gemineoa* (B; note the difference in scale between species). Predictions are shown for 2050 and 2100 under low (RCP45) and high (RCP85) CO_2_ emission scenarios. Points are locations from which individuals were sampled. Note that species have planktonic life stages (gametes, embryos, and larvae) that spend days to weeks offshore before transitioning to sessile life stages (juveniles and adults) onshore in the intertidal.

Other cross-validations further supported the robustness of our results. Patterns in Figure 2 were similar to those from an equivalent analysis that included Moran’s eigenvector maps to account for population structure in *G. gemineoa* (the only species for which genetic isolation was marginally associated with distance), with many overlapping SNPs (Table S5).

#### Genomic vulnerability to future climate change

Gradient forest models identified temperature-driven turnover in allele frequencies at 11 candidate loci in *G. caespitosa* (from 24 candidates screened), and 10 candidate loci in *G. gemineoa* (from 17 candidates screened) (Figure S5). Based on the relative contributions of temperature variables to predicted turnover (indicated by the lengths and alignments of vectors with biplot axes in Figure 4; see also importance values in Figure S6), maximum temperature and temperature noise structure are most important to turnover in *G. caespitosa*, whereas mean temperature and monthly temperature range are most important to turnover in *G. gemineoa*. Importance aside, the close alignments of their vectors in both biplots suggest that temperature range and noise structure otherwise act similarly on turnover in both species (Figure 4).

Mapping current genetic turnover for both species identified divergent patterns of adaptive genetic composition along the open coast (Figure 4). For *G. caespitosa*, predicted allele frequencies were relatively homogeneous (Figure 4B), apart from turnover between western populations and the sole northeastern sample mapping to changes in maximum and mean temperature (higher in bluer and purple regions respectively). For *G. gemineoa* predicted allele frequencies were more heterogeneous (Figure 4D), with turnover between western populations and (mostly) allopatric ones in the northeast mapping to changes in temperature range and mean (higher in greener and bluer regions respectively). For both species, marked turnover between the open coast and enclosed bays mapped to changes in temperature range and noise structure (both higher within bays).

Not surprisingly, more extreme warming (RCP45 *versus* RCP85 and 2050 *versus* 2100 scenarios) is predicted to increase divergence between current and future genetic compositions, and hence the genetic offset needed to maintain thermal adaptation, in both species (Figure 5). For *G. caespitosa*, greatest offset is predicted for enclosed bays and the northeast coast, unless extreme warming to 2100 demands adaptation throughout its range (Figure 5A). For *G. gemineoa*, genetic offset is more than double the magnitude predicted for *G. caespitosa*, and is greatest for western (sympatric) populations — again, unless extreme warming to 2100 demands adaptation throughout its range (Figure 5B).

## Discussion

With climate change redistributing biodiversity around the globe (Pecl et al., 2017), predicting species’ responses to future climates entails understanding their current ranges, whether they harbour or share genetic variants involved in climate adaptation, and how variants are distributed across landscapes and seascapes undergoing climate change. We set out to assess the distribution of neutral and adaptive genomic variation in sister foundation species — the marine tubeworms *G. gemineoa* and *G. caespitosa* — across a sentinel region for climate impacts. We found that species hybridize little despite uncovering sympatry in their ranges, harbour mainly species-specific variants involved in adaptation to differing components of temperature, and face different risks of maladaptation under projected changes in temperature. These results offer new insights into the potential disruption of evolutionary adaptation and species distributions by near-future climate change in coastal ecosystems.

Detection of sympatry in sister *Galeolaria* species overturns previous molecular support for limited overlap in their ranges (Halt et al., 2009; Styan et al., 2008), but accords with their capacity for long-distance dispersal in early life (Olsen et al., 2020; Palumbi, 1994). Notably, the extent of range overlap detected here far exceeds estimates of poleward range shifts by marine species in the hotspot during the last decade (Sunday et al., 2015). Our results could therefore reflect more intensive sequencing across the hotspot here than in previous work. Moreover, that species show little gene flow in sympatry suggests the presence of strong reproductive barriers between them, despite maintaining a reasonable capacity to cross-fertilise and produce viable larvae (Styan et al., 2008). It is therefore possible that species in sympatry remain isolated by genetic incompatibilities arising at later postzygotic stages (Fierst & Hansen, 2010; Sinervo & Calsbeek, 2003), or other mechanisms (e.g., asynchronous gamete release, conspecific sperm precedence; Howard, 1999; Lotterhos & Levitan, 2010) that avoid hybridisation in the first place, and such possibilities warrant further research. Last, neutral genomic variation revealed low levels of population differentiation and moderate levels of inbreeding in both species, as seems to be common for external fertilisers with long-distance dispersal and limited control of mate choice (Olsen et al., 2020; Palumbi, 1994). However, other measures of neutral diversity were lower in *G. gemineoa* than *G caespitosa*, suggesting that species-specific reductions in population size may have left one species more genetically depauperate, and hence more vulnerable to decline, than the other (Reed & Frankham, 2003; Sgrò et al., 2011).

Climate adaptation also seems to differ between *Galeoalaria* species, given that putatively-adaptive loci show species-specific associations with different components of temperature — specifically, with its maximum and stochasticity for *G. caespitosa*, but its mean and range for *G. gemineoa*. Australia’s east coast is characterized by clear latitudinal gradients in the annual mean and seasonality of temperature driven by seasonal cycling of the East Australian Current, whereas the south coast is characterised by less-structured changes in temperature occurring longitudinally (Frusher et al., 2014; Waters, 2008). Hence, adaptive genetic variation in *G. gemineoa* (whose range extends northward along the east coast) associates most strongly with the dominant components of temperature variation throughout its range, as does adaptive variation in *G. caespitosa* (whose range is largely restricted to the south coast). This result emphasises the expected coupling of physical processes (e.g., oceanographic forcing) and evolutionary processes in the sea (Lotterhos et al., 2021), also detected in the handful of studies to so far link adaptive variants to physical characteristics of coastal ecosystems (Nielsen et al., 2021; Vranken et al., 2021; Wood et al., 2021). It may further suggest that sister *Galeoalaria* species have adapted to different selective pressures mediated by different components of temperature variation, facilitating poleward range shifts in *G. gemineoa*, especially, if conditions to which it has already adapted on the east coast extend southward with ongoing climate change.

Another possibility is that *Galeolaria* species have adapted to similar components of temperature variation, but differ in the genetic basis of adaptation in ways that affect power to detect associations between adaptive variants and those components. Supporting this idea, adaptation in both species is polygenic and involves loci that not only associate with different components of temperature to different degrees within species, but also overlap little between species. On one hand, this could reflect the multidimensional nature of climatic variables (Garcia et al., 2014; Waldock et al., 2018) if their different components drive selection at different genomic regions. Future studies could therefore assess whether putatively-adaptive loci are functionally associated with different traits that aid adaptation (e.g., Popovic & Riginos 2020), for example, to changes in mean temperature *versus* stochasticity in temperature. On the other hand, genetic differentiation of populations and species across the southeast hotspot is often attributed to their historical isolation during glacial maxima (Dawson, 2005; O’Hara & Poore, 2000). Consequently, *Galeolaria* species may have diverged genetically long before adapting to contemporary climates, and the relative contributions of isolation and adaptation to divergence in these (and other) lineages across the hotspot are currently hard to elucidate (Miller et al., 2013; Waters, 2008). Climate adaptation is nonetheless cited as a key driver of divergence in other local species (Miller et al., 2020; Wood et al., 2021). If such is also the case for *Galeolaria*, then barriers between sister species could be maintained by genetic incompatibilities arising from divergent adaptation (Dettman et al., 2007; Keller & Seehausen, 2012), in addition to the neutral divergence noted above. The nature and origin of these barriers, however, remains to be tested.

Mapping genomic vulnerability to future climate change across the hotspot predicts that *G. gemineoa* is substantially more vulnerable than *G. caespitosa*, generally needing twice as much genomic change to track climate change via evolutionary adaptation (Capblancq et al., 2020; Hoffmann et al., 2021). This may be due *G. gemineoa*’s distribution across a broad thermal gradient in the hotspot, leading to greater breadth of adaptation, and hence greater potential for future loss of adaptation in this species compared to *G. caespitosa*. Vulnerability also varied geographically within species, with higher levels predicted for populations approaching the northern edge of *G. caespitosa*’s range, and those approaching the western edge of *G. gemineoa*’s range (though we cannot rule out its extension further west than sampled). Compared to populations at range cores, range-edge populations often harbour novel genetic variants that can facilitate adaptation, but may also have higher risks of decline due to smaller population sizes and lower genetic diversity (Eckert et al., 2008; Polechová & Barton, 2015; Sexton et al., 2009). Our predictions of vulnerability may therefore flag potential range contractions in both species under future climate change, with *G. gemineoa* at comparatively greater risk due to its lower genetic diversity noted above. Last, *G. caespitosa* is predicted to have relatively high vulnerability in enclosed bays, marked by less flow and more extreme temperatures compared to open coasts (Barton et al. 2012). Hence, the relatively strong signals of climate adaptation detected in bays, shown here to harbour different adaptive variants to those found on nearby coasts, may also be prone to disruption under future climate change.

Despite their promise for inferring climate adaptation (and its predicted loss) in non-model organisms with limited tractability to experimentation (Fitzpatrick et al., 2021), the genomic tools used here have limitations that should be acknowledged (reviewed in Capblancq et al., 2020; Hoffmann et al., 2021; Rellstab et al., 2021). For instance, predictions of genomic vulnerability do not account for the ability of populations to adapt to climate change using standing genetic variation, or gene flow from other, well-adapted populations across a species’ range (which, as noted, could especially benefit *G. gemineoa*). Genotype-environment associations are also inherently correlative, and can be prone to false positives (Hoban et al., 2016; Rellstab et al., 2015; Tiffin & Ross-Ibarra, 2014), although we cross-validated adaptive candidates using multiple approaches here.

Notwithstanding such limitations, using genotype-environment associations to predict genomic vulnerability may point out populations requiring greater adaptation to track future climates, given that inherent costs of adaptation are expected to impose demographic pressure on populations while they adapt to projected changes (Bell, 2012; Haldane, 1957). For any focal organism, future studies should ideally aim to link putatively-adaptive genetic variants to variation in individual phenotypes and fitness, in addition to population growth and adaptive capacity, in order to improve and validate predictions of genomic vulnerability under climate change.

Overall, we present new insights into climate adaptation, its predicted disruption by climate change, and the implications for partly sympatric foundation species that enhance biodiversity in a sentinel region for climate change impacts. Identifying so-called evolutionarily significant units worth conserving for their genetic uniqueness, adaptive significance, and risk of decline, is one of the most pressing challenges facing us today, and a necessary step in developing proactive conservation strategies (Foden et al., 2019; Smith et al., 2014; Willi et al., 2022). Our findings advance that goal by identifying sister *Galeolaria* species as lineages on distinct adaptive trajectories linked to climate, that seemingly share little gene flow (and hence little scope to gain neutral diversity or climate-adaptive variants from one another), and are predicted to fare differently in future climates. As foundation species, moreover, future changes in either of their distributions will likely cascade to broader impacts on the biological communities they sustain (Thomsen et al., 2022). In this context, studies such as ours could enhance the holistic assessment of species vulnerability to climate change (Hoffmann et al., 2015; Williams et al., 2008), and contribute to the evolutionarily enlightened management of biodiversity in coastal ecosystems.

## Acknowledgements

We thank Chris Lee for valuable help with laboratory work, Javiera Olivares for valuable help in sampling of specimens, and Fisheries Victoria and Parks Victoria for collection permits. This research was supported by a Holsworth Wildlife Research Endowment awarded to CG, and by grants awarded under the Australian Research Council’s Discovery Scheme to KM and KH.

