## Supplemental materials for "Climate adaptation and vulnerability of foundation species in a global change hotspot"

**Supplementary materials**

Table S1. Sampled locations and their geographic coordinates, ordered from west to northeast (see Figure 1).

| Location | Latitude | Longitude |
|---|---|---|
| Glenaire | -38.7826 | 143.4254 |
| Cape Otway | -38.8542 | 143.5485 |
| Apollo Bay | -38.7615 | 143.6799 |
| Kennett River | -38.6698 | 143.8646 |
| Lorne | -38.5477 | 143.9869 |
| Anglesea | -38.4283 | 144.1827 |
| Torquay | -38.3396 | 144.3278 |
| Queenscliff | -38.2675 | 144.6672 |
| Dimmicks Beach | -38.3832 | 144.7773 |
| Flinders | -38.4756 | 145.0257 |
| YCW Beach | -38.5046 | 145.2514 |
| Cape Woolamai | -38.5305 | 145.3425 |
| Kilcunda | -38.5533 | 145.4787 |
| Eagles Nest | -38.6694 | 145.6708 |
| Walkerville | -38.8606 | 146.0006 |
| Picnic Bay | -39.0144 | 146.2912 |
| Norman Beach | -39.0349 | 146.3111 |
| Sealers Cove | -39.0201 | 146.444 |
| Yanakie | -38.8203 | 146.2666 |
| Red Bluff | -37.8676 | 148.0623 |
| Salmon Rocks | -37.8094 | 148.7262 |
| Py-yoot Bay | -37.7887 | 148.8863 |
| Clinton Rocks | -37.7788 | 149.1958 |
| Fly Cove | -37.7561 | 149.4972 |
| Pebbly Beach | -37.6121 | 149.7196 |
| Mallacoota | -37.571 | 149.7641 |
| Cape Howe | -37.5164 | 149.9624 |
| Greenglades | -37.2824 | 149.9429 |
| Eden | -37.0622 | 149.9102 |
| Merimbula | -36.8909 | 149.9298 |

Table S2. Temperature variables explored in this study, calculated for sampled locations using high-resolution (1 $km^2$ grid cell) daily observations of sea surface temperature from January 2010 to December 2018. Variables are categorised by components of change in temperature, with those used for analyses in bold. Names in brackets indicate calculations based on the WorldClim scheme (Fick and Hijmans 2017; Hijmans et al. 2005). Noise structure ($\beta$, capturing autocorrelation between daily observations) was calculated as the negative slope of the line, $log_{10}$ *spectral density* = $log_{10}$ *frequency*, fitted to observations in each cell according to Vasseur and Yodzis (2004). Noise structure is completely uncorrelated if $\beta = 0$ and positively correlated if $\beta > 1$).

| Component of change | Temperature variable |
|---|---|
| *Trend* | **Mean (BIO1)** |
| *Variability* | **Mean monthly range (BIO2)** |
| | Mean annual range (BIO7) |
| | Isothermality (BIO3) |
| | Temperature seasonality (BIO4) |
| *Extremity* | **Maximum temperature of warmest month (BIO5)** |
| | Minimum temperature of coldest month (BIO6) |
| | Mean temperature of warmest quarter (BIO10) |
| | Mean temperature of coldest quarter (BIO11) |
| *Stochasticity* | **Noise structure ($\beta$)** |

Table S3. Estimates of genetic diversity (averaged across loci) for populations of *G. caespitosa* and *G. gemineoa*. N is the number of individuals analysed, Ho is their observed heterozygosity, $H_S$ is their expected heterozygosity, $F_{IS}$ is their inbreeding coefficient, and AR is their allelic richness (AR).

| Population | N | $H_O$ | $H_S$ | $F_{IS}$ | AR |
|---|---|---|---|---|---|
| ***Galeolaria caespitosa*** | | | | | |
| **Glenaire** | 3 | 0.060 | 0.092 | 0.213 | 1.173 |
| **Cape Otway** | 9 | 0.066 | 0.101 | 0.249 | 1.329 |
| **Apollo Bay** | 8 | 0.067 | 0.096 | 0.202 | 1.299 |
| **Kennett River** | 9 | 0.070 | 0.101 | 0.238 | 1.335 |
| **Lorne** | 6 | 0.072 | 0.100 | 0.203 | 1.276 |
| **Anglesea** | 9 | 0.071 | 0.099 | 0.203 | 1.340 |
| **Torquay** | 10 | 0.068 | 0.099 | 0.232 | 1.348 |
| **Queenscliff** | 4 | 0.072 | 0.094 | 0.119 | 1.202 |
| **Dimmicks Beach** | 8 | 0.072 | 0.101 | 0.207 | 1.323 |
| **Flinders** | 8 | 0.070 | 0.102 | 0.223 | 1.325 |
| **YCW Beach** | 8 | 0.060 | 0.100 | 0.301 | 1.319 |
| **Cape Woolamai** | 7 | 0.070 | 0.104 | 0.220 | 1.313 |
| **Kilcunda** | 4 | 0.064 | 0.100 | 0.218 | 1.216 |
| **Eagles Nest** | 6 | 0.065 | 0.101 | 0.242 | 1.278 |
| **Walkerville** | 10 | 0.069 | 0.097 | 0.213 | 1.358 |
| **Picnic Bay** | 4 | 0.064 | 0.100 | 0.248 | 1.217 |
| **Norman Beach** | 4 | 0.070 | 0.097 | 0.177 | 1.225 |
| **Sealers Cove** | 5 | 0.081 | 0.102 | 0.136 | 1.275 |
| **Yanakie** | 8 | 0.069 | 0.095 | 0.195 | 1.312 |
| **Pebbly Beach** | 1 | 0.073 | - | - | 1.073 |
| ***Galeolaria gemineoa*** | | | | | |
| **Glenaire** | 10 | 0.056 | 0.085 | 0.270 | 1.291 |
| **Apollo Bay** | 1 | 0.063 | - | - | 1.063 |
| **Kennett River** | 2 | 0.062 | 0.081 | 0.122 | 1.125 |
| **Anglesea** | 1 | 0.063 | - | - | 1.063 |
| **Torquay** | 1 | 0.039 | - | - | 1.039 |
| **Queenscliff** | 5 | 0.062 | 0.082 | 0.161 | 1.219 |
| **Cape Woolamai** | 7 | 0.054 | 0.087 | 0.270 | 1.259 |
| **Kilcunda** | 1 | 0.058 | - | - | 1.058 |
| **Picnic Bay** | 3 | 0.044 | 0.073 | 0.268 | 1.120 |
| **Norman Beach** | 5 | 0.062 | 0.089 | 0.189 | 1.212 |
| **Sealers Cove** | 10 | 0.055 | 0.087 | 0.266 | 1.305 |
| **Red Bluff** | 7 | 0.058 | 0.085 | 0.218 | 1.241 |
| **Salmon Rocks** | 9 | 0.066 | 0.086 | 0.171 | 1.296 |
| **Py-yoot Bay** | 10 | 0.054 | 0.081 | 0.244 | 1.288 |
| **Clinton Rocks** | 10 | 0.059 | 0.083 | 0.215 | 1.293 |
| **Fly Cove** | 9 | 0.056 | 0.080 | 0.216 | 1.261 |
| **Pebbly Beach** | 9 | 0.067 | 0.085 | 0.159 | 1.305 |
| **Mallacoota** | 10 | 0.058 | 0.079 | 0.189 | 1.298 |
| **Cape Howe** | 10 | 0.061 | 0.091 | 0.252 | 1.326 |
| **Greenglades** | 8 | 0.061 | 0.085 | 0.199 | 1.281 |
| **Eden** | 7 | 0.055 | 0.089 | 0.262 | 1.273 |
| **Merimbula** | 6 | 0.054 | 0.085 | 0.267 | 1.219 |

Table S4. Associations between temperature variables and candidate loci identified for *G. caespitosa* and *G. gemineoa* by redundancy analysis (RDA) *versus* BayPass analysis. For redundancy analyses, numbers of loci with association strengths (correlations with temperature variables) > |0.5| are shown in brackets.

| **Species** | **Temperature variable** | **Method** | | | | **Overlap between methods** |
|---|---|---|---|---|---|---|
| | | **RDA** | **BayPass** | | | |
| | | | XtX outliers | Significant associations | Both | |
| *G. caespitosa* | Mean | 181 (26) | 1,471 | 29 | 5 | 0 |
| | Maximum | 230 (44) | | 37 | 19 | 10 |
| | Mean monthly range | 213 (34) | | 30 | 15 | 6 |
| | Noise structure | 151 (38) | | 30 | 19 | 5 |
| *G. gemineoa* | Mean | 248 (14) | 1,459 | 26 | 16 | 4 |
| | Maximum | 93 (38) | | 15 | 9 | 2 |
| | Mean monthly range | 173 (37) | | 9 | 2 | 2 |
| | Noise structure | 165 (50) | | 11 | 5 | 0 |

Table S5. Associations between temperature variables and candidate loci identified for *G. gemineoa* by redundancy analysis with distance-based Moran's eigenvector maps included to account for population structure. This was the only species for which genetic isolation was marginally associated with geographic distance. The model and all four ordination axes were statistically significant ($P < 0.001$), consistent with the main analysis without eigenvector maps included (see Figure 2). Numbers of loci with association strengths (correlations with temperature variables) > |0.5| are shown in brackets.

| **Temperature variable** | **Candidate loci** | **Overlap with main analysis** |
|---|---|---|
| Mean | 188 (9) | 88 |
| Maximum | 105 (22) | 61 |
| Mean monthly range | 213 (32) | 83 |
| Noise structure | 186 (31) | 75 |

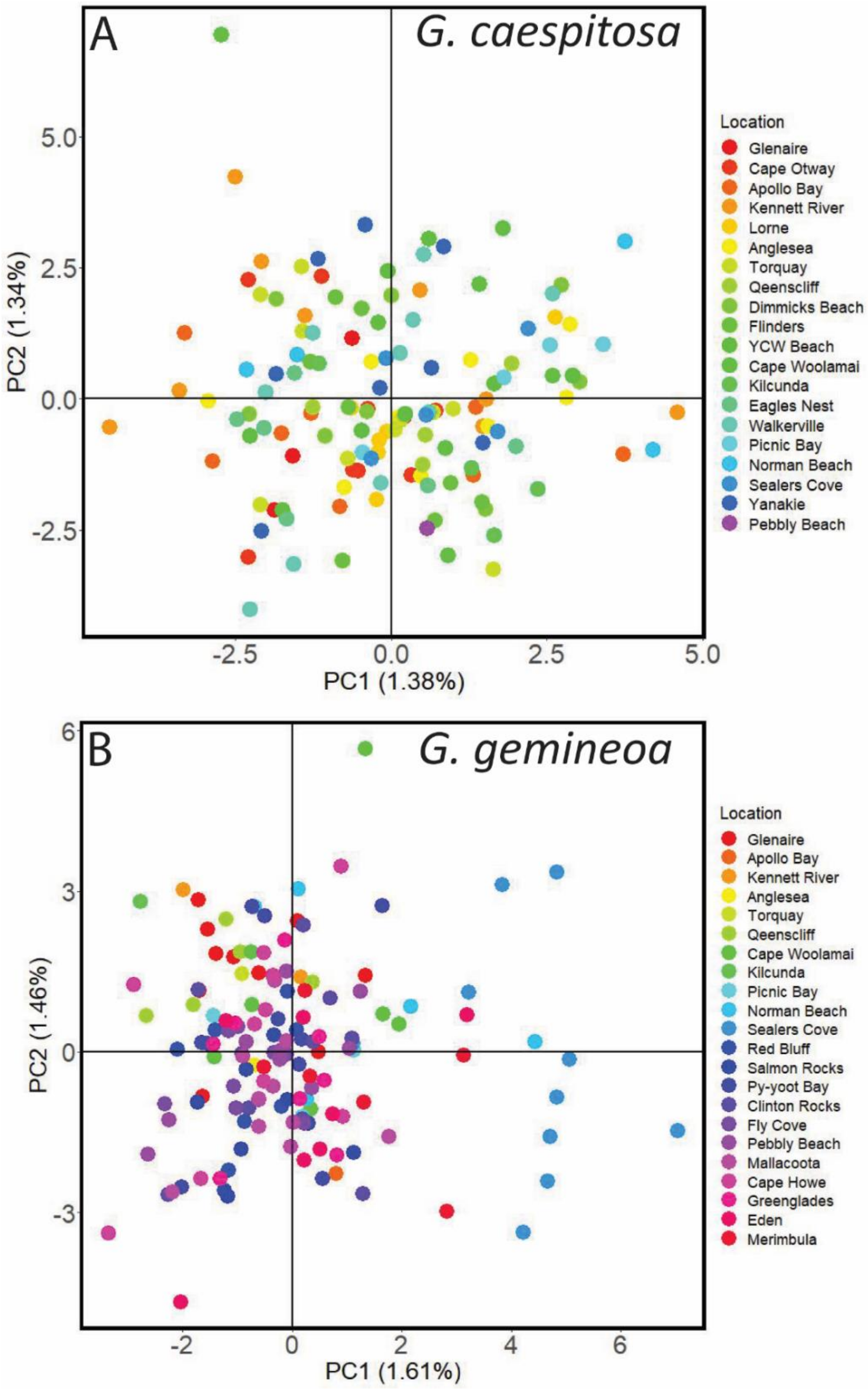


Figure S1. Lack of population structure detected by principal component analyses of genetic variation performed separately for (A) *G. caespitosa* and (B) *G. gemineoa*.

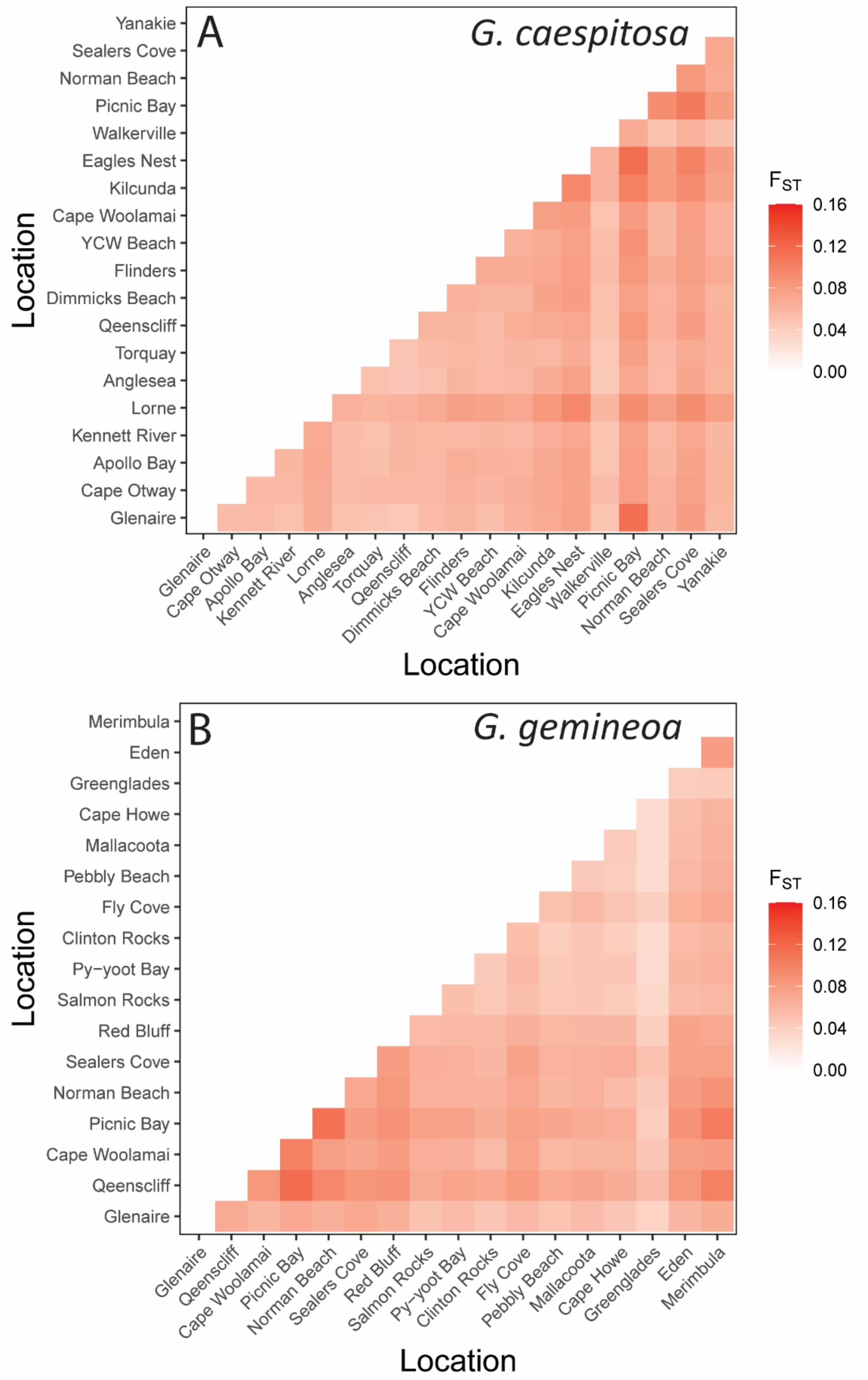


Figure S2. Pairwise genetic distances ($F_{ST}$) between populations of (A) *G. caespitosa* and (B) *G. gemineoa* sampled from different locations. Sampled locations are ordered from west to northeast (with geographic distance increasing from bottom to top and from left to right).

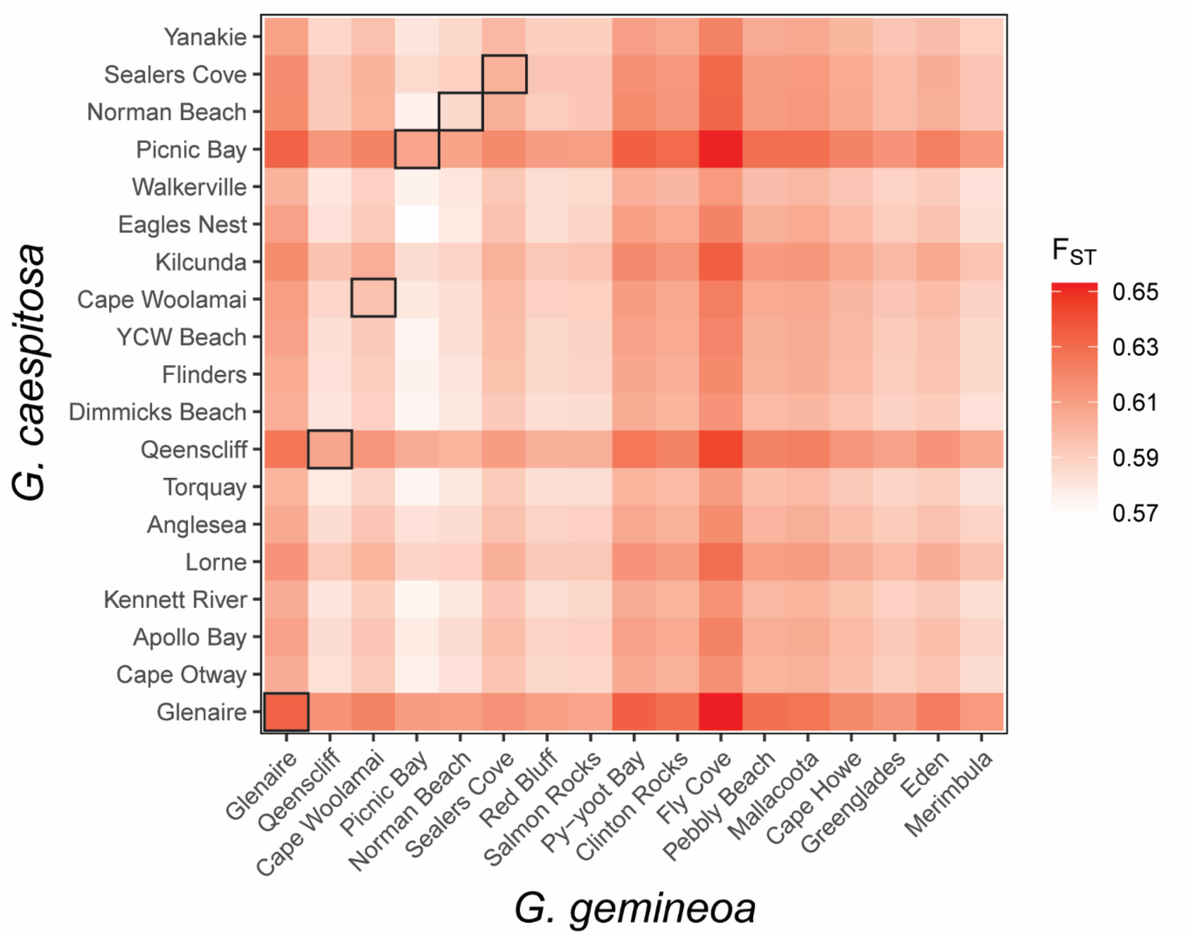


Figure S3. Pairwise genetic distances ($F_{ST}$) between species, with sympatric populations in black squares and allopatric populations in open squares. Sampled locations are ordered from west to northeast (with geographic distance increasing from bottom to top and from left to right).

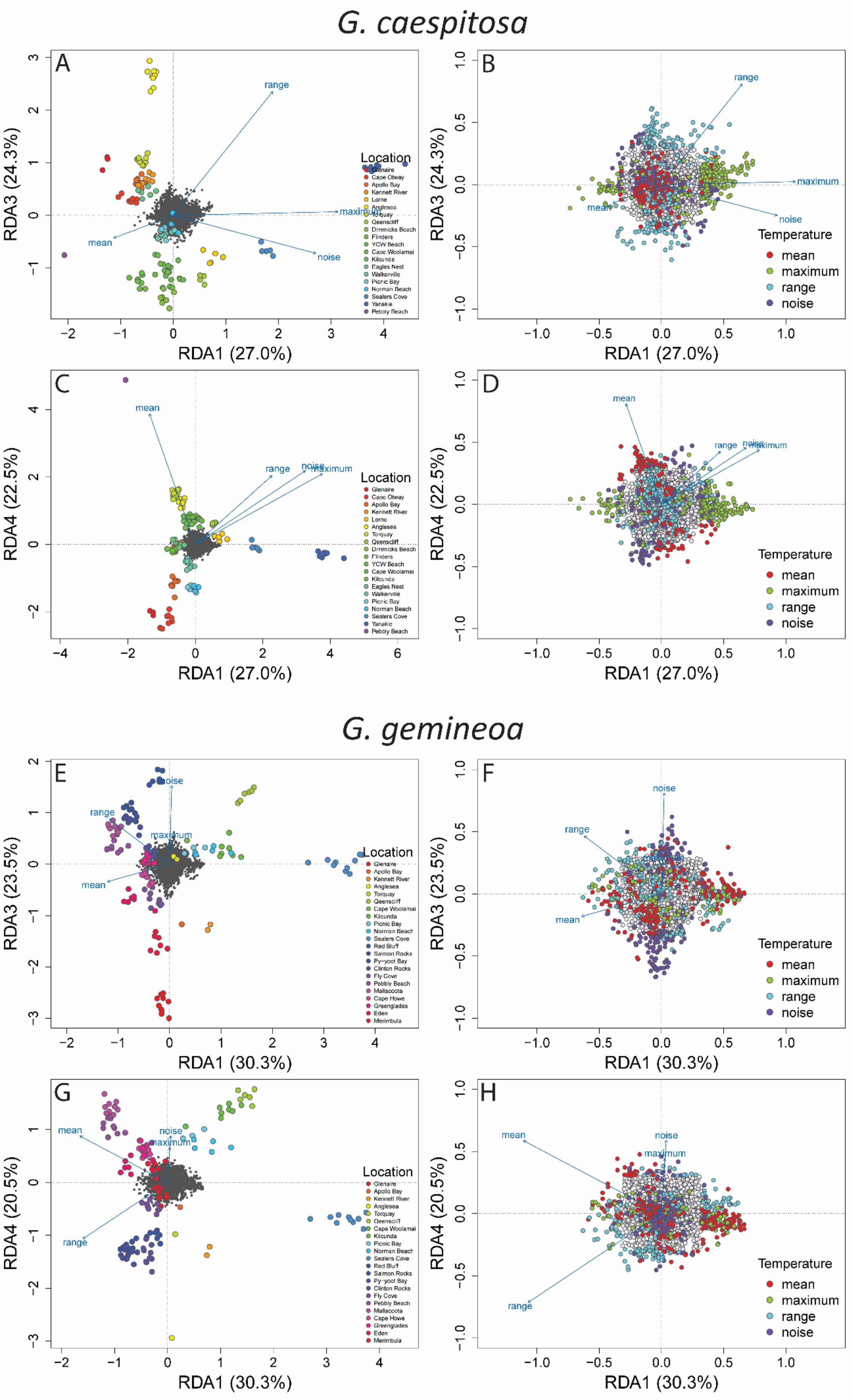
G. caespitosa
A
RDA3 (24.3%)
RDA1 (27.0%)
range
maximum
noise
mean
Location
Glenaire
Cape Otway
Apollo Bay
Kennett River
Lorne
Anglesea
Torquay
Qeenscliff
Dimmicks Beach
Flinders
YCW Beach
Cape Woolamai
Kilcunda
Eagles Nest
Walkerville
Picnic Bay
Norman Beach
Sealers Cove
Yanakie
Pebbly Beach
B
RDA3 (24.3%)
RDA1 (27.0%)
Temperature
mean
maximum
range
noise
C
RDA4 (22.5%)
RDA1 (27.0%)
D
RDA4 (22.5%)
RDA1 (27.0%)
G. gemineoa
E
RDA3 (23.5%)
RDA1 (30.3%)
Location
Glenaire
Apollo Bay
Kennett River
Anglesea
Torquay
Qeenscliff
Cape Woolamai
Kilcunda
Picnic Bay
Norman Beach
Sealers Cove
Red Bluff
Salmon Rocks
Py-yoot Bay
Clinton Rocks
Fly Cove
Pebbly Beach
Mallacoota
Cape Howe
Greenglades
Eden
Merimbula
F
RDA3 (23.5%)
RDA1 (30.3%)
G
RDA4 (20.5%)
RDA1 (30.3%)
H
RDA4 (20.5%)
RDA1 (30.3%)

Figure S4. Associations between genotype and temperature variables identified for *G. caespitosa* (A–D) and *G. gemineoa* (E–H) by redundancy analysis. Biplots show ordination axes (RDA1 *versus* RDA3, and RDA1 *versus* RDA4) explaining associations not presented in Figure 2. In all panels, closer alignments of items with ordination axes indicate stronger associations with axes. In (A), (C), (E,) and (G), grey points are single loci, other points are individuals coloured by location, and vectors are variables. In (B), (D), (F), and (H), which magnify left-hand plots to focus on loci, candidate loci (identified as significant outliers on ordination axes) are coloured by the variable they associate most strongly with.

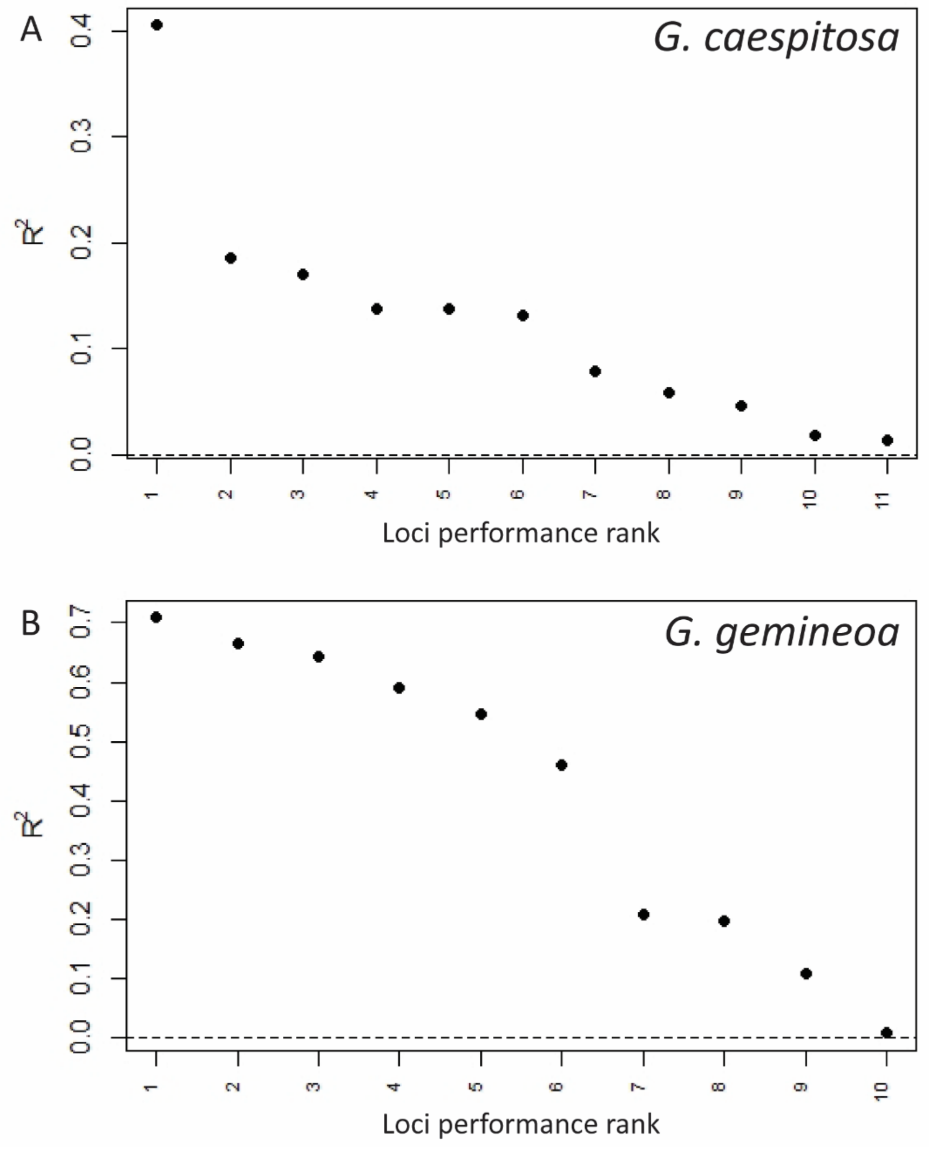


Figure S5. $R^2$ measure of the fit of the gradient forest model for *G. caespitosa* (A) and *G. gemineoa* (B) for each retained candidate loci.

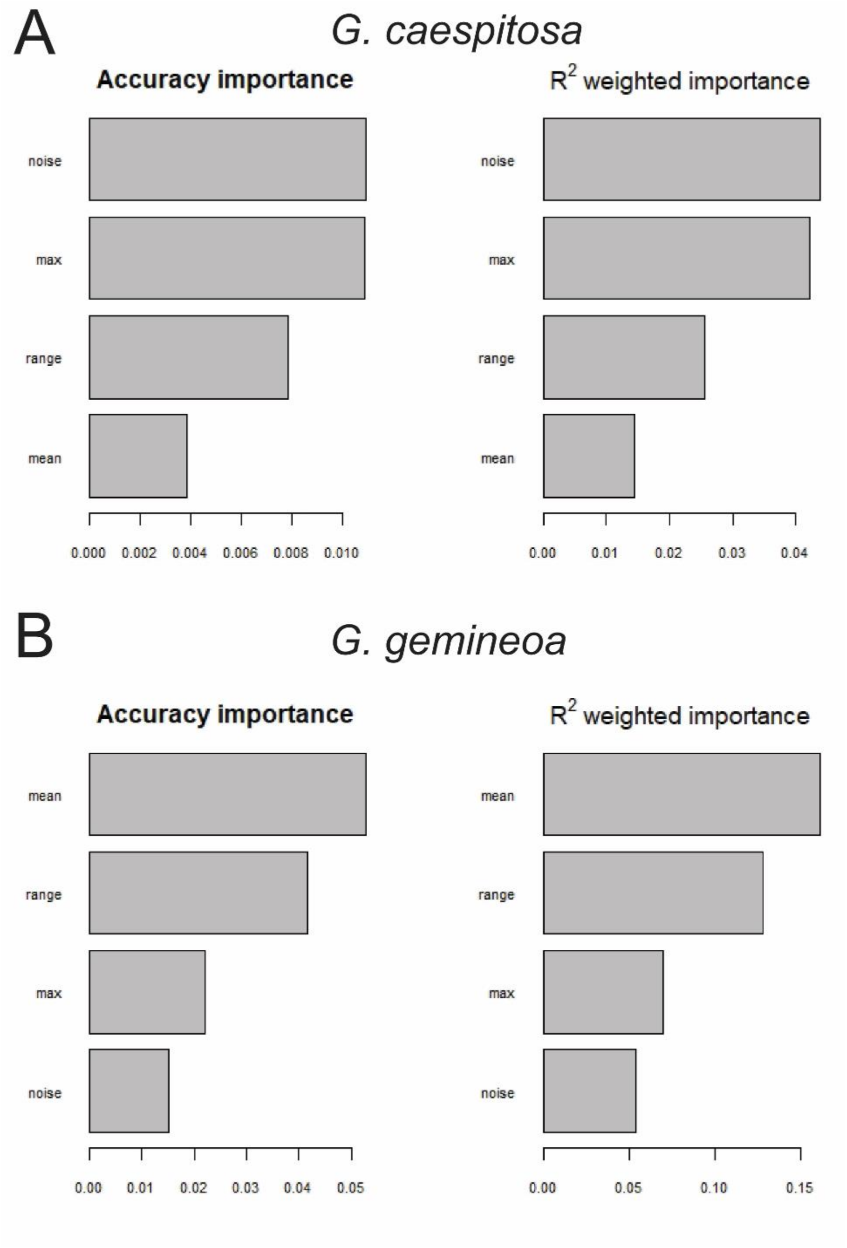


Figure S6. Summaries of gradient forest models for *G. caespitosa* (A) and *G. gemineoa* (B). Accuracy importance averaged across loci (left) and average importance weighted by loci $R^2$ (right). Both measure the relative contributions of temperature variables to predicted turnover in allele frequencies at candidate loci included in each model.
